# Coarse-grained Modeling and Experimental Investigation of the Viscoelasticity of Human Gut Mucus and Nanoparticle Dynamics

**DOI:** 10.1101/2023.02.28.530485

**Authors:** Liming Zhao, Sandra L. Arias, Ilana L. Brito, Jingjie Yeo

## Abstract

A thick layer of mucus covering the gastrointestinal tract acts as an innate barrier guarding the epithelial surface. The high molecular weight and cross-linked glycoproteins (mucins), the major building blocks of mucus, can effectively obstruct or trap invading noxious substances, such as detrimental bacteria and virus. The mucus layer as well as any trapped material can be regularly removed by the friction force from food flow and gastrointestinal peristalsis, the process of which primarily relies on the viscoelastic and shear-thinning properties. Conversely, the process by which beneficial substances, such as drug nanoparticles, cross the mucus layer and contact the epithelium is also influenced by the chemical and rheological properties of the mucus layer. Gastrointestinal disorders, most notably colitis, are often accompanied by changes to the mucosal structure. In this study, we experimentally characterized the viscoelasticity and dynamic viscosity of mucus collected from human intestinal cells. In addition, we developed a bi-component mesoscopic-scale mucus model that contained Muc2, the dominant mucin secreted in healthy individuals, and Muc5AC, which is secreted by intestinal goblet cells in certain intestinal disorders. This model enabled us to study the effects of cross-linking and mucin concentration on rheological properties of mucus. Furthermore, we quantified changes in the diffusion dynamics of nanoparticles in mucus networks caused by factors such as the size of nanoparticles, nanoparticle-mucin interactions, and the degree of mucin cross-linking.

## Introduction

The gastrointestinal (GI) tract is coated by a thick layer of mucus hydrated gel, which is primarily composed of water (95%), mucin glycoproteins (5%), and minor substances such as electrolytes, lipids, nucleic acids, etc.^1,2^ Human mucins, the main building blocks of the gel, are a large family that have more than twenty unique genes.^3^ Among these, Muc2 is the predominant mucin secreted within the intestinal tract in healthy individuals. During inflammatory bowel disease and colorectal cancer, epithelial cells begin to express Muc5AC and Muc5B, in addition to Muc2^4,5^. Mucins are primarily secreted by goblet cells.^6,7^ Muc2 and Muc5AC have similar chemical structures that consist of a protein backbone containing more than 5,000 amino acids and hydrophilic glycan side chains. The protein backbone is partitioned: hydrophobic N-terminal and C-terminal are located at two ends, with hydrophobic cysteine domains and glycosylated regions in the middle.^8^

The main functions of the mucus layer are to lubricate and protect the epithelium from mechanical stress, exposure to luminal contents and the entry of pathogenic microorganisms, including bacteria and viruses.^9,10^ The lubricating and protective functions of the mucus layer depend primarily on its mechanical properties, which are affected by several factors, including the rate of mucus secretion and erosion caused by frictional forces during peristalsis, as well as mucus degradation derived from microbial activities.^11^ Besides its barrier functions, the mucus layer can also significantly limit the effectiveness of drug delivery systems. Alterations of the GI mucus layer are associated with several gastrointestinal syndromes, including acute intestinal infections, ulcerative colitis, Crohn’s disease, and colorectal cancer.^12,13^

Some efforts have been put on simulating mucus over the past ten years. Gao *et al*. used molecular dynamics (MD) simulations to investigate the interaction between the mucins and the nanomaterials, but the dynamics information in the mesoscopic scale could not be obtained due to the size limitation of the full-atomistic MD simulation.^14^ Even though other studies developed mesoscopic-scale mucus model, the cross-linking network of mucus in some studies were too idealized^15,16^, and others only modeled single-component and monodisperse mucins^17–19^. These studies did not consider the diversity of mucins and the complexity of the mucus network. Thus, developing more complex models that allow detailed mechanistic studies of mucus’ mechanical behavior and chemicals’ dynamics is a critical step toward understanding the mucus barrier *in vivo*, predicting the pharmacokinetics of nanoparticles and nanocarriers, and optimizing DNA- and phage-based therapies. In this study, we developed mesoscopic-scale models specifically for human gut mucins. We calibrated the model by comparing the geometry (length, diameter, pore size, *etc*.) and the mechanical properties of mucins to experimental measurements obtained from human cells and published values. Based on the model, we studied the diffusion dynamics of polyethylene glycol (PEG) nanoparticles inside of the mucus.

## Materials and Methods

### Mucus collection

The human colorectal adenoma cell line HT29-MTX-E12, which adopts a goblet-like phenotype, was cultured in 75 cm^2^ plastic flasks in Dulbecco’s Modified Eagle’s Medium (DMEM) supplemented with 10% fetal bovine serum (FBS) for 21 days after confluence. At day 21, the serum-containing medium was replaced by advanced DMEM/F12 (Gibco) without serum to reduce mucus contamination by foreign proteins. At day 22, the serum-free culture medium was discarded, and the flasks were gently rinsed with Dulbecco’s phosphate-buffered saline (DPBS) several times. After that, the cells were incubated for 45 min at 37 °C in DPBS with 0.5 mM forskolin to release the mucus layer. The mucus covering the cells was then gently removed in DPBS (with calcium and magnesium, pH=7.4) and used immediately.

### Mechanical characterization of pristine mucus

A rotational rheometer DHR-3 (TA instruments) was used to investigate the viscoelastic characteristics and apparent viscosity of freshly isolated mucus by applying frequency and flow sweeps. First, the linear viscoelastic region was determined using an amplitude sweep at a constant frequency of 1 rad/s using a 40 mm parallel plate geometry. The mucus’ viscoelastic properties in terms of loss (G”) and storage (G’) moduli were then determined using frequency sweeps performed between 0.01 to 100 rad/s at a 0.5 uNm constant torque with 5 to 10 measurement points per decade and 300 um separation gaps. The mucus’ apparent viscosity was measured using flow sweeps in steady mode with a cone and plate geometry (40 mm cone diameter, 63 um truncation). The samples were subjected to shear rates between 0.1 to 100 s^-1^, with a 300 s equilibration time and 300 um separation gaps. Amplitude, frequency, and flow sweeps were all performed in DPBS with calcium and magnesium (pH=7-7.2) at 25 °C using three different cell culture batches and at least two samples per batch.

### Coarse-grained model of mucins and NPs

We built a coarse-grained (CG) model of mucins in which mucin chains are represented by the bead-spring model. The Muc2 and Muc5AC molecules consist of hydrophobic protein beads (type *h*), which represent the N-terminals, C-terminals and cysteine-rich domains, and hydrophilic beads (type *g*), which represent the glycosylated regions. We estimated the location and range of the N-terminal, C-terminal, and cysteine-rich domains based on the amino acid sequences of the Muc2 (UniProt: Q02817) and Muc5AC (UniProt: T1S9D5).^8^ As a result, the ratio and the position of type *h* beads for each mucin can be adjusted in accordance with their respective amino acid sequences. PEG nanoparticles were type *p* beads packed into spheres. The structures of Muc2, Muc5AC and nanoparticles are shown in Figure 1. We also added water beads (type *w*), such that the mucins and NPs are able to diffuse in an aqueous solution.

**Figure 1.**
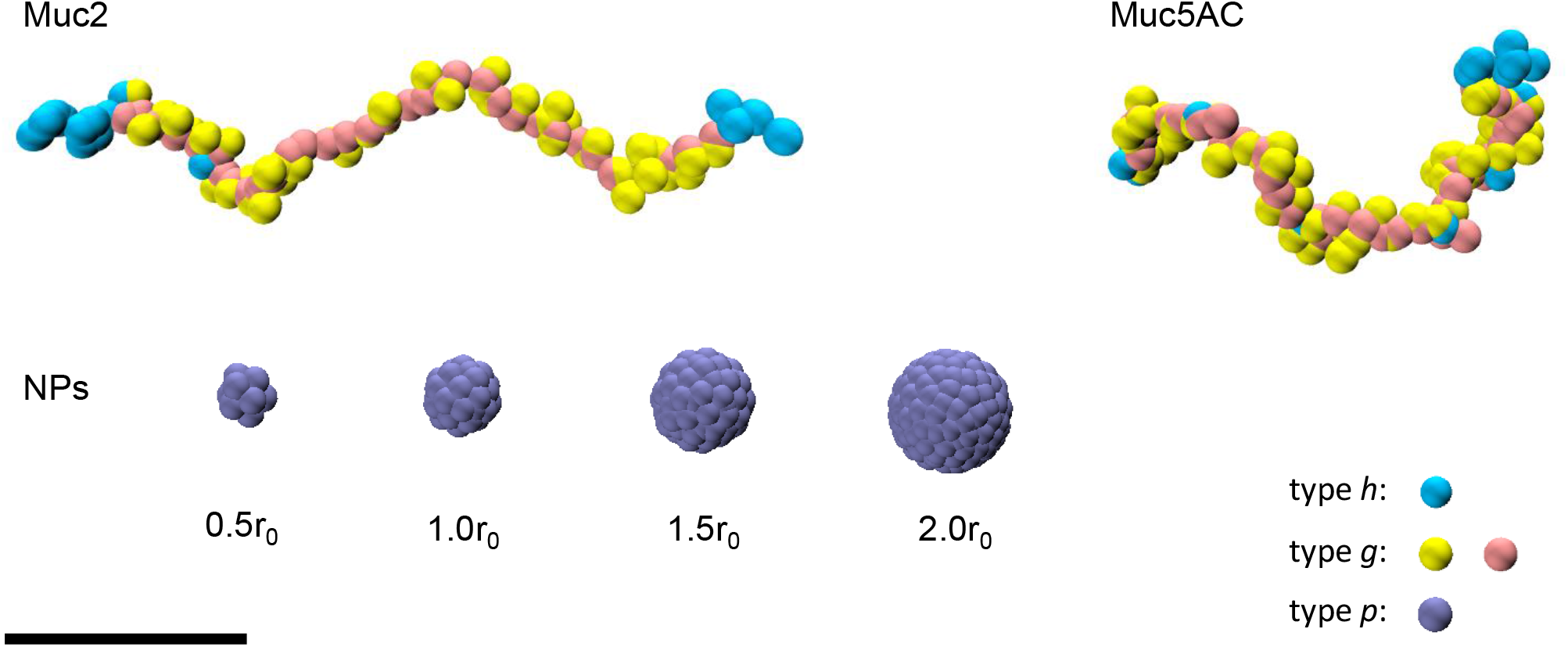
CG model structures of Muc2, Muc5AC and nanoparticles. The cyan beads represent N-terminal, C-terminal, and cysteine-rich domains. The yellow and pink beads represent the glycosylation regions. The iceblue beads represent nanoparticles. The scalebar denotes 10 length units (10 r_0_).

The dimensions of the mucin molecules in the simulation were determined according to the previous literature.^19–23^ First, the number of beads in the mucin backbone were polydisperse, which obeyed a Gaussian distribution with a mean of 50 beads and a standard deviation of 3. The length distribution was proportional to the measurement results (671 nm in average within the range from 579 nm to 752 nm).^20^ An average of two beads made up each side chain, resulting in the ratio of the diameter to the mucin length comparable with the observation from the microscope and the previous simulation model.^19,21^ Additionally, hydrophilic beads accounted for 87% of the total mucins in our model, which was close to the mass proportion of 80% or 90% mentioned in the literature.^22,23^

### Force field of the CG model

To model the interactions between beads, we adopted the interaction potential from the dissipative particles dynamics (DPD) method, as it captures the hydrophilic-hydrophobic relationship in detail between different types of beads.^24^ The interaction forces in the DPD model within the cut-off radius *r*_*c*_ are defined as the sum of the conservative force *F*_*ij*_^*C*^, dissipative force

*F*_*ij*_^*R*^, and random force *F*_*ij*_^*R*^:

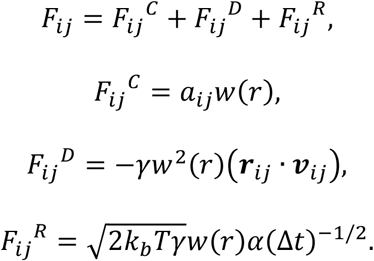

As DPD uses normalized non-dimensional units, the cut-off radius *r*_*c*_ was set to one unit length (*r*_*c*_ = *r*_0_) and the temperature was 1 *k*_*b*_*T*. The values of the parameters *a*_*ij*_ used in our model are listed in Table 1. The repulsive parameters between PEG and hydrophobic protein beads *a*_*p*ℎ_ and between PEG and glycan beads *a*_*pg*_ were calculated from Hansen solubility parameters^25^:

**Table 1.**
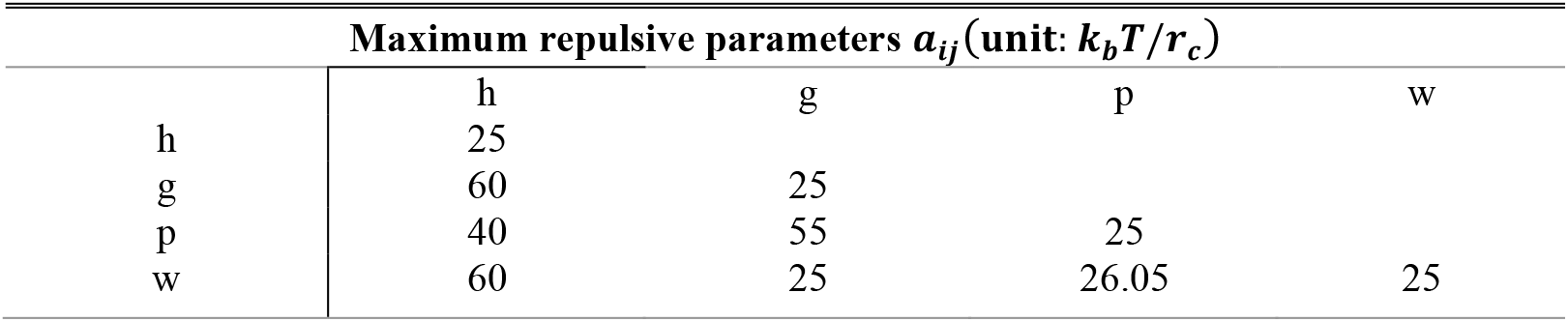
The maximum repulsive parameters *a*_*ij*_ used in the DPD model.

| Maximum repulsive parameters $a_{ij}$ (unit: $k_b T / r_c$) | | | | |
| --- | --- | --- | --- | --- |
|  | h | g | p | w |
| h | 25 |  |  |  |
| g | 60 | 25 |  |  |
| p | 40 | 55 | 25 |  |
| w | 60 | 25 | 26.05 | 25 |

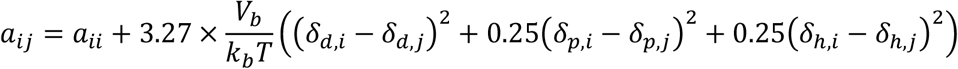

where *a*_*ii*_ is the self repulsive parameter (25 *k*_*b*_*T*/*r*_0_ in our model) and *V*_*b*_ is the volume of a bead. Due to a lack of published solubility parameters (*δ*_*d*_, *δ*_*p*_, *δ*_ℎ_) of the glycan and protein in mucins, their *a*_*ij*_ parameters were estimated using the solubility of representative glycan units of sucrose and dextran to calculate the repulsive parameter between PEG and the glycan in mucins (*a*_*pg*_ = 55 *k*_*b*_*T*/*r*_0_).^25^ Similarly, using zein proteins, we generalized the solubility parameters to obtain the repulsive parameter between PEG and the protein in mucins (*a*_*p*ℎ_= 40 *k*_*b*_*T*/*r*_0_).^25^ The parameter between PEG and water *a*_*pw*_ was adopted from Luo *et al*.,^26^ and other parameters *a*_*ij*_ were adopted from Moreno *et al*.^19^ The scalar *w*(*r*) solely depends on the distance between two particles. The parameter for the dissipative force *γ* was set to 4.5. The variable *α* is a Gaussian noise.^24^

In addition, we also defined the bond and the angle potentials as follows. The adjacent beads in a backbone or in a side chain were connected by harmonic bonds: *U*_*b*_=*K*_*b*_(*r*_*ij*_−*r*_0,*b*_)^2^, where the stiffness *K*_*b*_ was set to 25 *k*_*b*_*T*, and the equilibrium distance *r*_0,*b*_ was set to 0.7 *r*_0_.^19^ The adjacent three *g* beads in the backbone were connected by a harmonic angle: *U*_*a*_=*K*_*a*_(*θ*_*ijk*_−*θ*_0,*a*_)^2^, where the stiffness was set to

### Equilibrium and cross-linking

We first randomly generated polydisperse mucin molecules in the simulation box and equilibrated them by running 600,000 steps of NVE ensemble followed by 10,000,000 steps of NVT ensemble (dt = 0.01). Then, if necessary, we extracted the coordinates of type *h* beads, and created intermolecular disulfide bonds if the distance between any two *h* beads was smaller than 2r_0_. In this study, we built mucus model with 0%, 25%, 50%, 75%, and 100% cross-linking, meaning that the corresponding proportion of disulfide bonds satisfying the criteria above were created. After adding disulfide bonds, we ran 10,000,000 steps of NVT ensemble to re-equilibrate the system.

### Pore size calculation

Pore size distribution was calculated by PSDsolv.^27,28^ For an arbitrary point in the simulation box, the pore size of the point is defined as the maximum radius of a sphere passing through the point, such that the sphere has no overlap with any beads. The accuracy was set to 0.2r_0_, *i*.*e*., the bin width of the pore size distribution was 0.2, and the error tolerance was 0.02.

### Hydrophobic node construction

Hydrophobic nodes were composed of type *h* beads. We defined a node as a cluster of type *h* beads in which all type *h* beads were separated at most by 2r_0_. Additionally, a node must have contained type *h* beads from at least two different mucin molecules. Thus, the nodes represent the regions where the cross-linking occurred. We used the Depth-First Search algorithm (DFS)^29^ to find the hydrophobic nodes and denoted them as single spheres in Figure 5.

### Steady shear simulation

The steady shear simulations were conducted in the NVT/SLLOD ensemble. To avoid processors losing bonds and avoid unrealistically high viscosities, bonds were broken (deleted) if the bond length was more than 2 times of the length in equilibrium. According to previous studies of protein (collagen) and chemical bonds, this threshold was determined to be the critical point for breaking such chemical bonds.^30,31^ The peak viscosity values were recorded during steady shear as the final viscosities.

### Nanoparticles Diffusion

Nanoparticles were randomly added in the empty spaces within the mucin network after the mucins were equilibrated. The radii of NPs were 0.5r_0_, 1.0r_0_, 1.5r_0_, and 2.0r_0_, and hence their volumes varied as well. We added NPs into the model systems such that the volume of NPs was 0.2 times of the volume of the mucins. The diffusions were conducted under the NVT ensembles for at least 1,000,000 steps (dt = 0.02) to ensure that the NPs were in the diffusive regime, rather than the ballistic regime. During the diffusion, each NP was treated as a rigid body, so every *p* beads in the NP had the same velocity.

## Results

### Mechanical and rheological properties of mucin

We conducted dynamic oscillatory shear experiments to investigate the flow behavior of mucus derived from HT29-MTX-E12 cells via dynamic rheological tests. Rheological measurements relate the viscoelastic properties of polymers to molecular structure and modes of molecular motion, as well as their dependence on molecular weight, molecular weight distribution, concentration, chemical structure, and other variables.^32^ Six measurements corresponding to different batches of *in vitro* cell culture were used to determine the storage (G’) and loss moduli (G”) of the mucus on the linear viscoelastic region (Figure 2). The G’ measures elastically recoverable deformation, whereas G” measures permanent deformation of the material. The mucus exhibited a viscoelastic response to deformation, with minimal change in both variables over an extensive range of frequencies. The values for the G’ were larger than the loss modulus, with values of 0.25 ± 0.24 Pa and 0.06 ± 0.04 Pa at one rad/s, respectively. This behavior is characteristic of highly elastic physical gels rather than a solution of entangled polymer chains, indicating that mucus derived from HT29-MTX-E12 cells forms a gel network.

**Figure 2.**
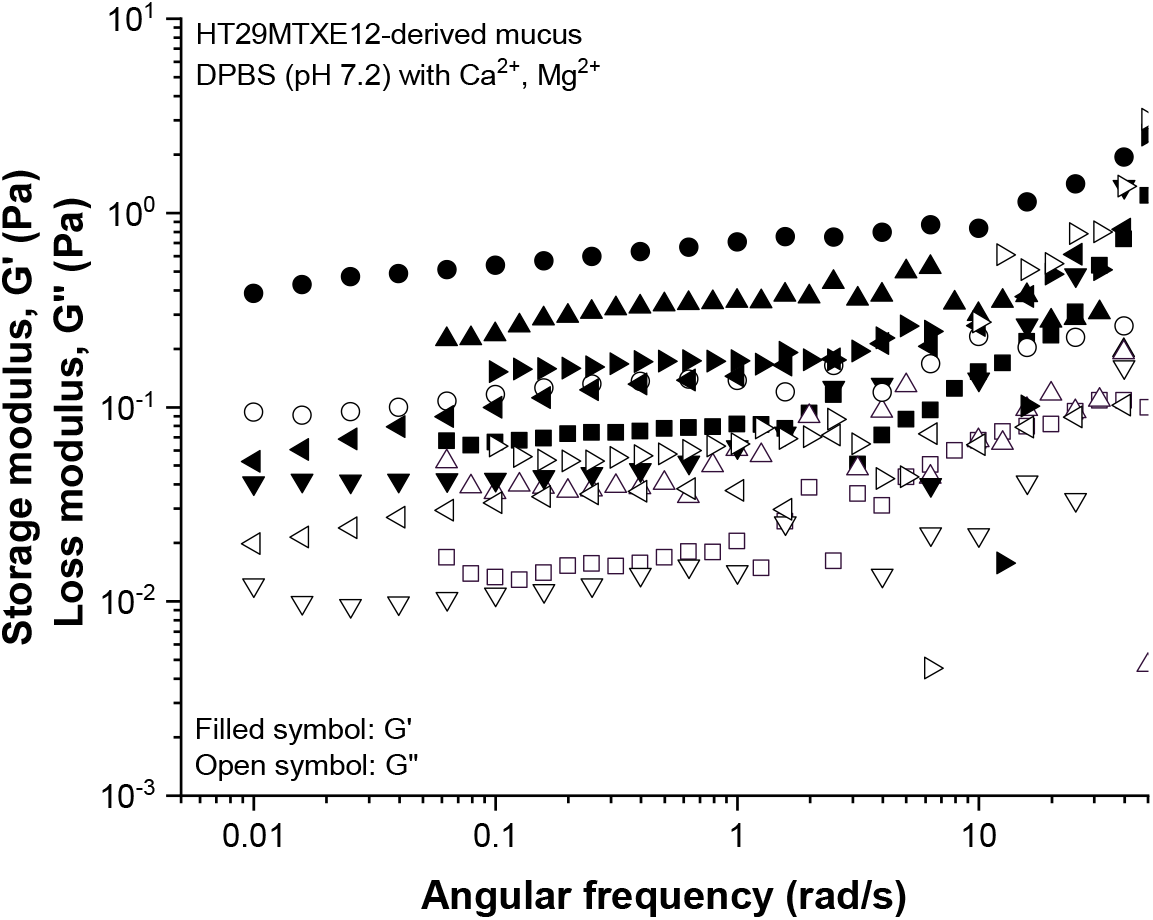
The storage (G^’^) and loss moduli (G^’’^) of six mucus samples collected from different batches of HT29-MTX-E12 cells.

The viscosity of mucus measured experimentally and extrapolated using our models under shear conditions agreed (Figure 3 (a)). The viscosity was exponentially decreased when the strain rate increased from 10^−1^ to 10^2^ s^-1^. To validate our DPD model, we calculated the viscosities of mucus by applying a steady shear to the simulation box. We obtained the viscosities of mucins with different degrees of cross-linking (Figure 3 (b)), the values of which were converted to SI units by scaling the values with the viscosity of pure water. The meaning and the controlling strategy of the degree of cross-linking were illustrated in the Methods section. Our results show that, as the degree of cross-linking increased from 0% cross-linking to 50% cross-linking, the viscosities increased dramatically. However, further increases in cross-linking density did not lead to further large increases in the viscosity, which plateaued instead. As we found that our mucus samples behaved more like elastic gels experimentally (Figure 2), we chose fully cross-linked (100%) mucin models to perform further simulations and characterizations. The viscosities of three simulation models with different mucin weight percentages were shown in Figure 3 (a). Due to limited temporal scales that could be captured in DPD simulations, the smallest applied strain rate was on the order of 10^5^ s^-1^, which was still higher than the maximum strain rate in the experiment. However, the extrapolated trend of the viscosity with increasing strain rates for 5 wt% mucins agreed with our experimental measurements.

**Figure 3.**
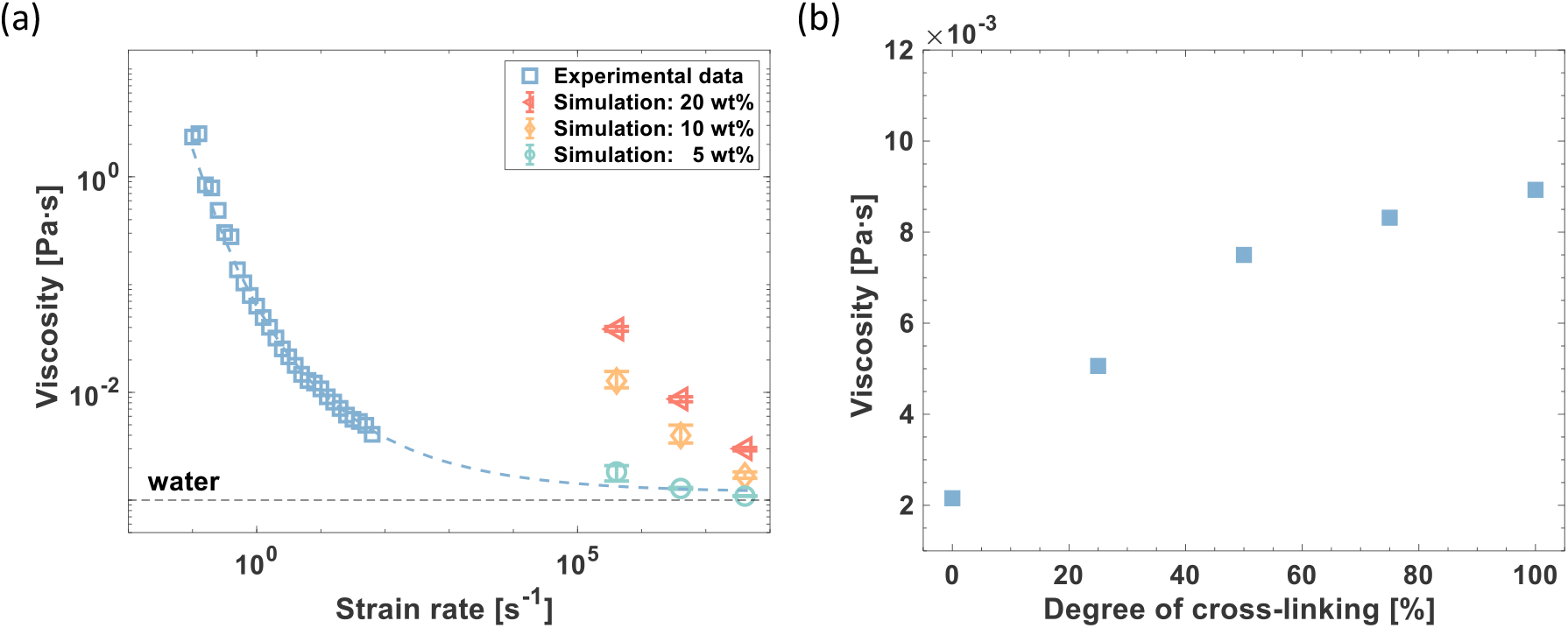
(a) The steady shear viscosities obtained from the experiments and simulation calculations. The blue dash line serves as the guide for the eyes. The viscosity of pure water, denoted by the black dash line, was given as the benchmark. (b) Simulation results of the viscosities of the mucins with 0%, 25%, 50%, 75%, and 100% degrees of cross-linking.

### Pore size measurement and aggregation characterization

Pore size is an important structural feature of polymer networks and affects the diffusion of nanoparticles or bacteria through mucus.^33–35^ Based on the storage modulus and the elastic blob theory^36,37^, the average pore size of mucus collected from the HT29-MTX-E12 cells was 247.92 ± 3.06 nm. In the simulation, we calculated the pore sizes of mucin under three different weight percentages, namely 5 wt%, 10 wt% and 20 wt%. In terms of the composition, mucus in the experiment was collected from HT29-MTX-E12 cells, which naturally produce mucus composed of 60% Muc5AC and 40% Muc2.^6,7^ Therefore, we used this Muc5AC/Muc2 ratio in the simulation and tested pure Muc5AC and pure Muc2 as a benchmark (Figure 4). We found that the average pore sizes were roughly proportional to the mucins weight percentage. For example, the mean pore sizes are 7.76r_0_, 4.69r_0_, and 2.36r_0_ (Muc5AC/Muc2= 0.6) when the weight percentages are of 5%, 10%, and 20%, respectively. In addition, the pore size decreases as the concentration of Muc5AC increases, which can be clearly observed in the 5 wt% simulation. To explain the changes in pore size, we examined the distributions of the hydrophobic nodes of mucins (Figure 5). At low weight percentages, Muc2 tended to phase-separate due to low numbers of crosslinks and aggregated to leave large empty spaces (Figure 5 (a)), but Muc5AC was more evenly distributed throughout the whole simulation domain (Figure 5 (c)). Compared with Muc2, Muc5AC has more cysteine-rich domains, denoted by hydrophobic beads (type *h*) in the simulation, and these domains are cross-linkable regions.^38–40^ Hence, crosslinking of these additional cysteine-rich domains helped the Muc5AC network maintain a more evenly distributed network structure with smaller pore sizes. This crosslinking effect became more evident for systems of pure Muc2 at 20 wt%: the hydrophobic nodes were evenly distributed throughout the simulation domain if there were sufficient numbers of crosslinks distributed throughout the network due to greater numbers of crosslink-able hydrophobic N-terminals and C-terminals (Figure 5 (d)). Thus, having more Muc5AC at 10 and 20 wt% did not reduce the average pore sizes to the same extent as in models with 5 wt% Muc2 because the pore sizes were already much smaller to begin with.

**Figure 4.**
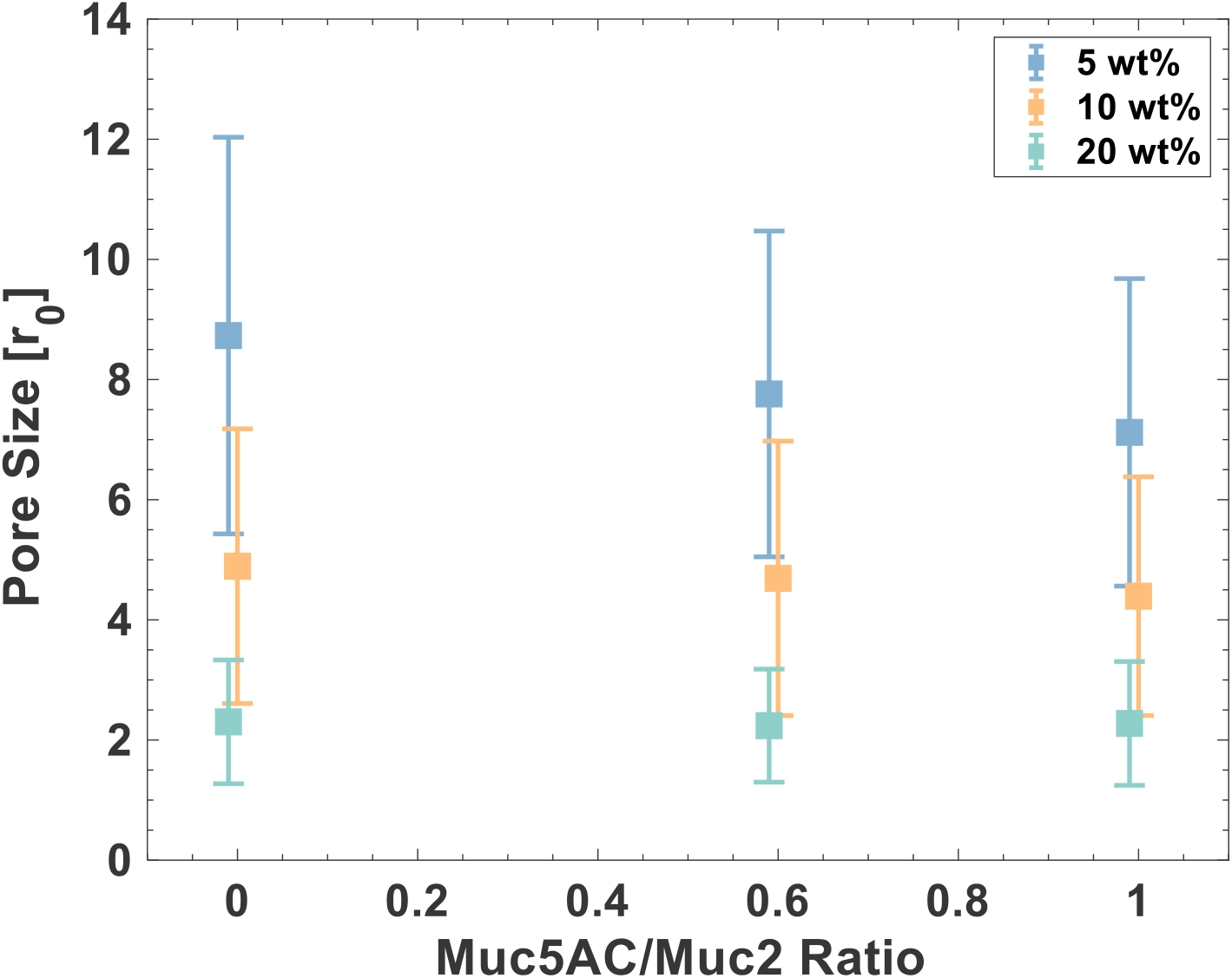
The pore sizes calculated in the simulations for different Muc5AC/Muc2 ratios and different weight percentages of total mucins, {0, 0.6, 1}×{5 wt%, 10 wt%, 20 wt%}. The x-coordinates were jittered for greater clarity of the error bars.

**Figure 5:**
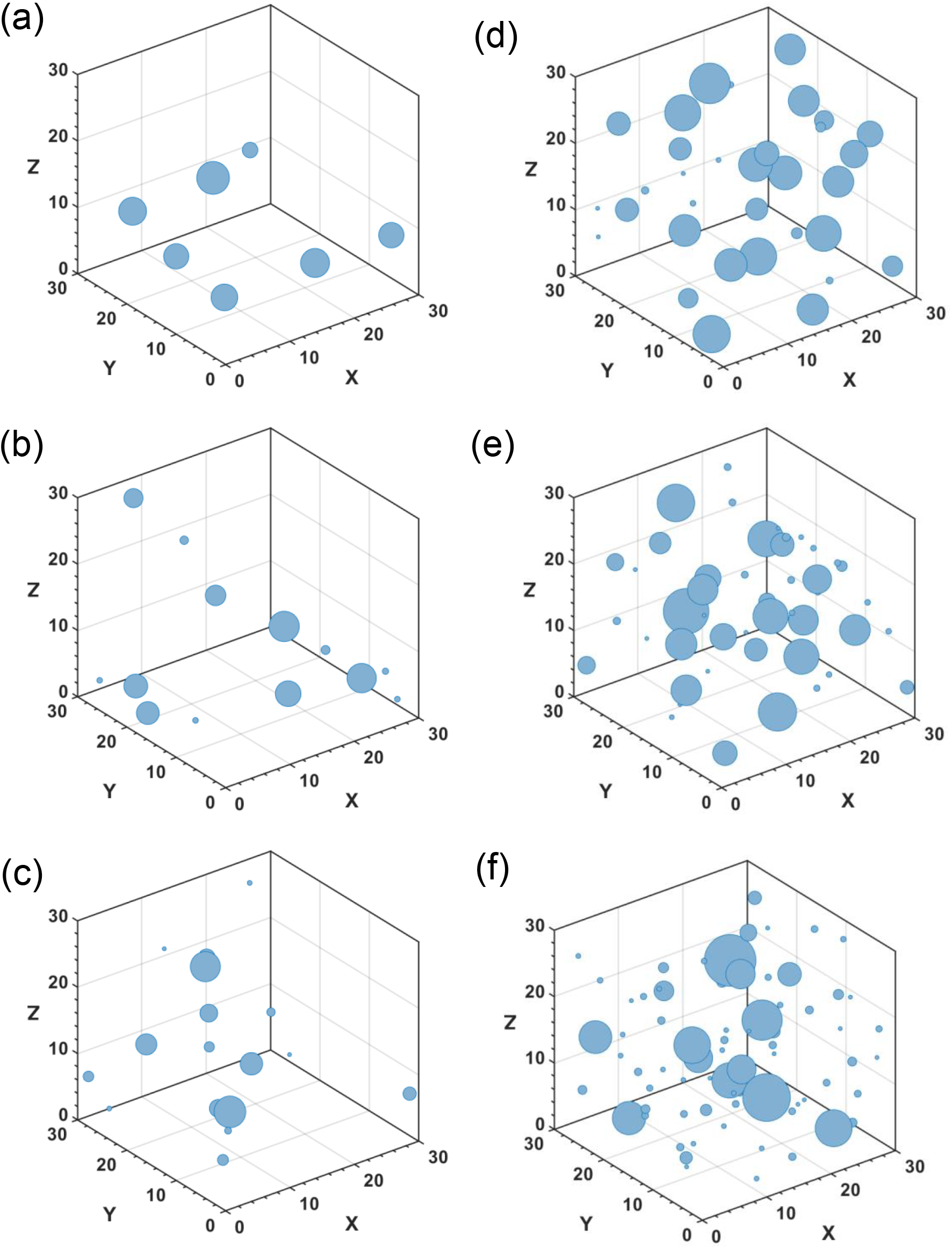
(a) – (c) The hydrophobic node distributions when the mucin is of 5 wt% and the ratio of Muc5AC/Muc2 is 0, 0.6, and 1, respectively. (d) – (f) The node distributions when the mucin is of 20 wt% and the ratio of Muc5AC/Muc2 is 0, 0.6, and 1, respectively. The sizes of the nodes indicate the number of beads in the nodes (range from 2 to 327).

### Nanoparticles dynamics

We examined the NP dynamics in mucus of NPs of different sizes. The range of NP sizes were chosen based on the time-averaged pore size distribution of mucins (Figure 7 (a)), which was normally distributed with a mean of 2.36r_0_. Our experimentally measured average pore size was 247.92 ± 3.06 nm, thus we added NPs with radii of 0.5r_0_, 1.0r_0_, 1.5r_0_, and 2.0r_0_ into our models. By considering the ratio of the NPs’ radii to the pore size of mucin, these NPs were representative of PEG NPs with radii of 50 nm, 100 nm, 150 nm, 200 nm, respectively. Also, this range of NP radii allowed us to investigate the NP dynamics when the NP radius was either much smaller than or comparable to the average pore sizes of mucins (Figure 6). Furthermore, these sizes conform to the actual radii (5 nm to 200 nm) used in NP-based therapeutics.^41–43^

**Figure 6:**
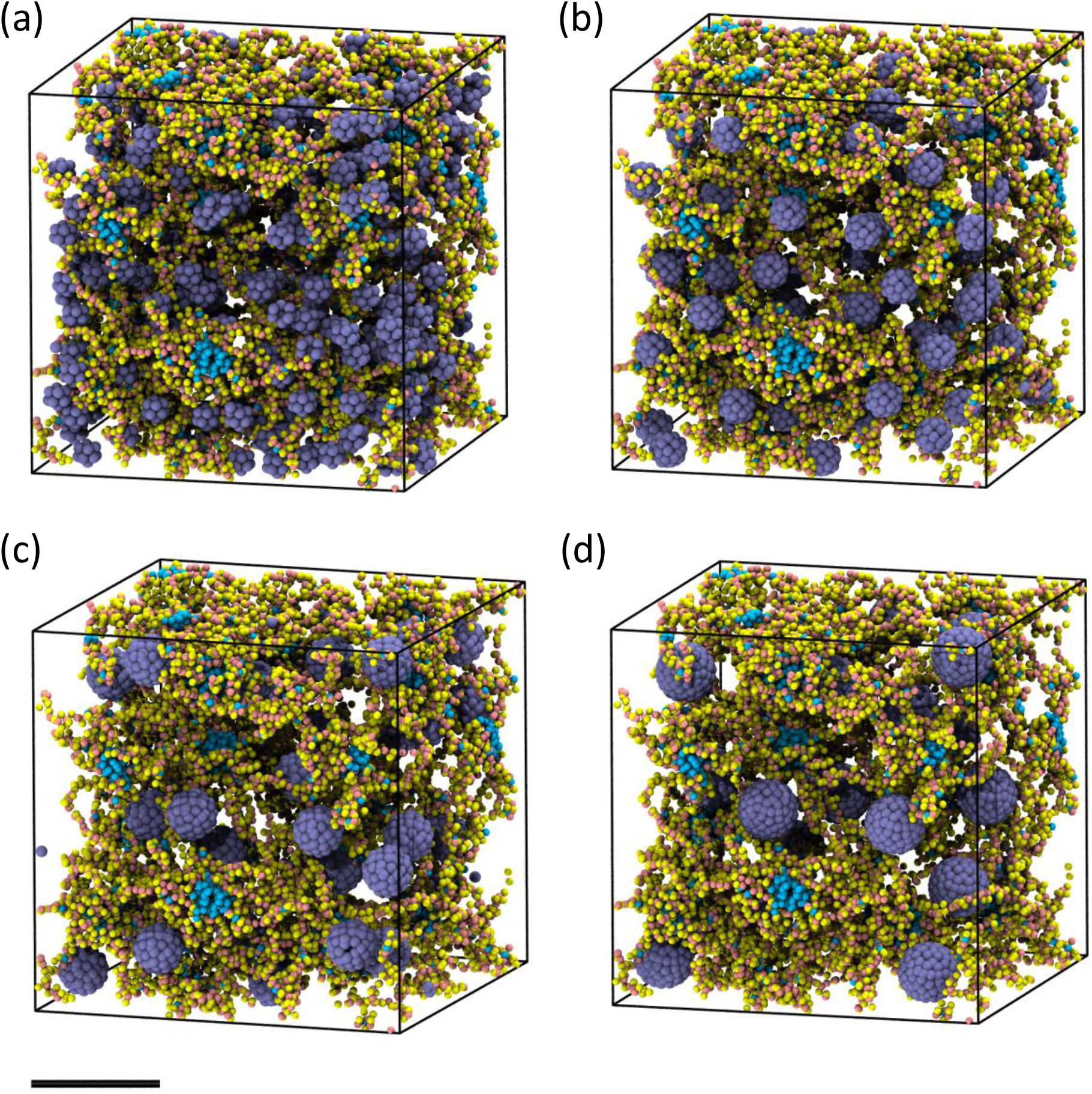
Snapshots of nanoparticles with different radii inside of 20 wt% mucins. The radii of NPs were (a) 0.5r_0_, (b) 1.0r_0_, (c) 1.5r_0_, and (d) 2.0r_0_. The weight percentages of NPs were same for all cases, namely one-fifth of the weight percentages of the mucins. The cyan beads represent N-terminal, C-terminal, and cysteine-rich domains. The yellow and pink beads represent the glycosylation regions. The iceblue beads represent nanoparticles. The scalebar indicates 10 DPD length units (10 r_0_).

**Figure 7:**
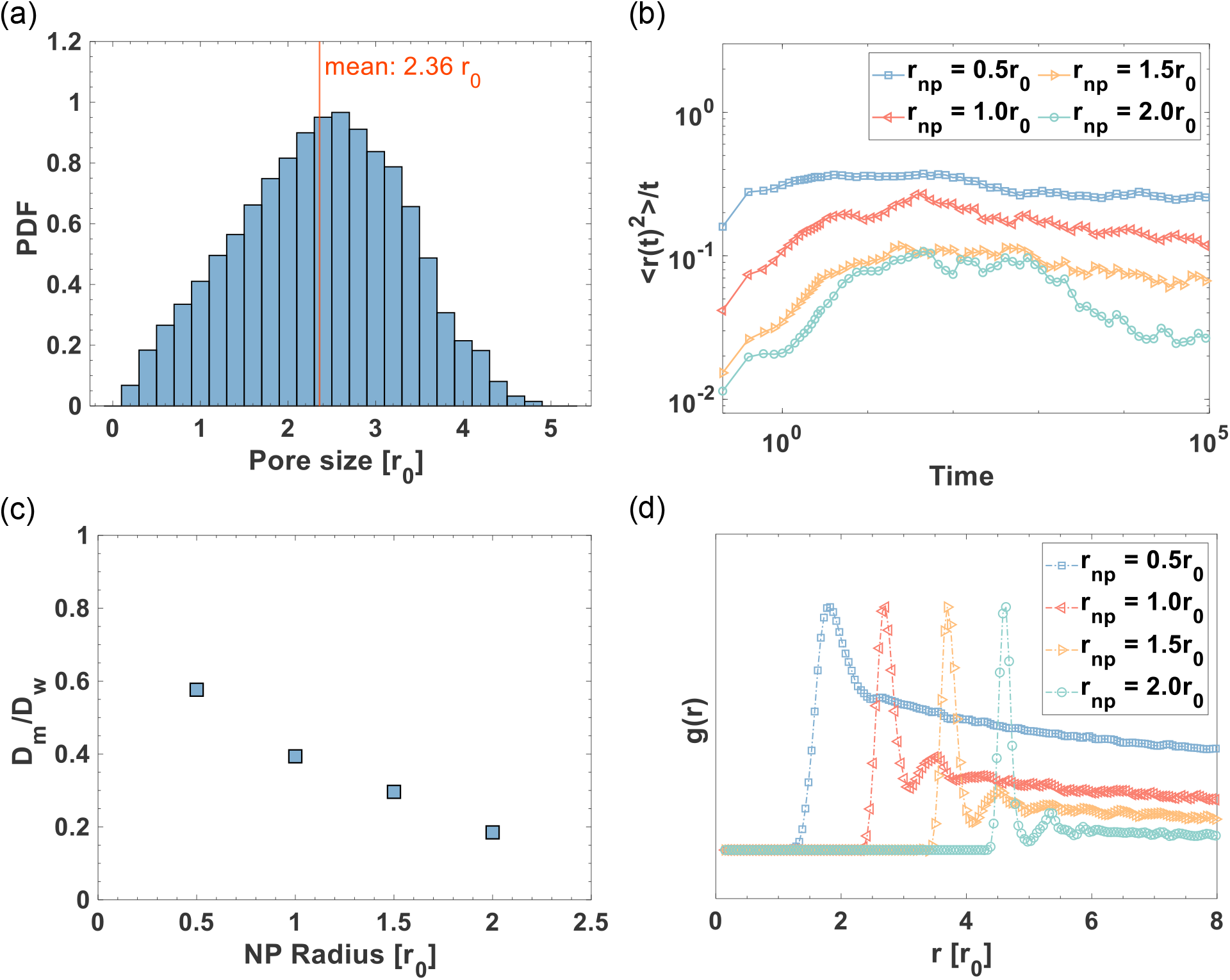
(a) The pore size distribution of the 20 wt% mucins after equilibrium. (b) The log-log plot of the mean squared displacements divided by time for NPs radii of 0.5r_0_, 1.0r_0,_ 1.5r_0_, and 2.0r_0_. (c) The ratios of the NP diffusion coefficients in mucus D_m_ to the diffusion coefficient in pure water D_w_ obtained from the diffusive regions in (b). (d) The NP-NP radial distribution functions of different sizes of NPs. The RDFs were normalized by dividing their maximum values.

The diffusion coefficient quantifies the dynamics of NPs and can be obtained, according to the Einstein relation, from the slope of the mean squared displacement (MSD) with respect to time.^44^ However, diffusion of nanoparticles in polymer networks usually goes through three distinct regimes: (1) the ballistic regime where the MSD is proportional to *t*^2^; (2) the subdiffusive regime where the MSD is proportional to *t*^*β*^ (*β* < 1); (3) the diffusive regime (Fickian diffusion) where the MSD is proportional to *t*.^45^ The diffusion coefficient describes the diffusion rate of nanoparticles within the diffusive regime. To clearly distinguish the three regimes, we analyzed the trend of MSD/time versus time (Figure 7 (b)). Initially, the NPs were in the ballistic regime where the curve sloped upwards with time. Large NPs (r = 2.0r_0_) underwent longer periods of ballistic diffusion than the small NPs (r = 0.5r_0_). Subsequently, the curves began to slope downwards, indicating that the NPs transitioned into the subdiffusive regime.

Eventually, all the curves plateaued when the NPs attained the diffusive regime and the diffusion coefficient was calculated. We further found that the weakening effect of the mucins on the NP diffusion rate became more significant with increasing NP size (Figure 7 (c)). To investigate the aggregation of NPs, we provided the normalized radial distribution functions (RDFs) for all four NPs (Figure 7 (d)). The highest peak (first peak) corresponded to the case in which NPs were closely aggregated and directly touching other NPs. The separation of the four highest peaks corresponded to the differences in NP radii. Except the small NPs, we could observe second peaks for the other three NPs, which implied that larger NPs tend to form aggregates with greater long-range order, but the small NPs tended to be distributed more evenly throughout the mucin network.

To ensure the robustness of our model parameters, we tested a series of *a*_*pg*_ and *a*_*p*ℎ_ and obtained their MSDs (Figure 8). As there were no previously reported maximum repulsive parameters between PEG and glycan *a*_*pg*_and between PEG and protein *a*_*p*ℎ_, as illustrated in the Method section, we obtained reference values of *a*_*pg*_and *a*_*p*ℎ_. It worth noting that *a*_*pg*_ primarily increased the kinetics during the ballistic regime, all plateaued to a similar range of average values during the diffusive regime (Figure 8 (a) and (c)). *a*_*p*ℎ_ neither significantly changed the ballistic regime nor the diffusive regime, which could be due to protein beads forming hydrophobic nodes that reduced the collision frequency with NPs and therefore had minimal impact on their diffusive motion (Figure 8 (b) and (d)). Therefore, we conclude that the diffusion rate is not sensitive to the parameters *a*_*pg*_ and *a*_*p*ℎ_. However, the parameters *a*_*pg*_ and *a*_*p*ℎ_ strongly influenced NPs aggregation. As shown in Figure 9 (a), larger *a*_*pg*_ induced higher repulsion between the glycan and SNPs, hence SNPs tended to aggregate more tightly. However, smaller and larger *a*_*pg*_ and *a*_*p*ℎ_ would facilitate the aggregation of LNPs (Figure 9 (c) and (d)).

**Figure 8:**
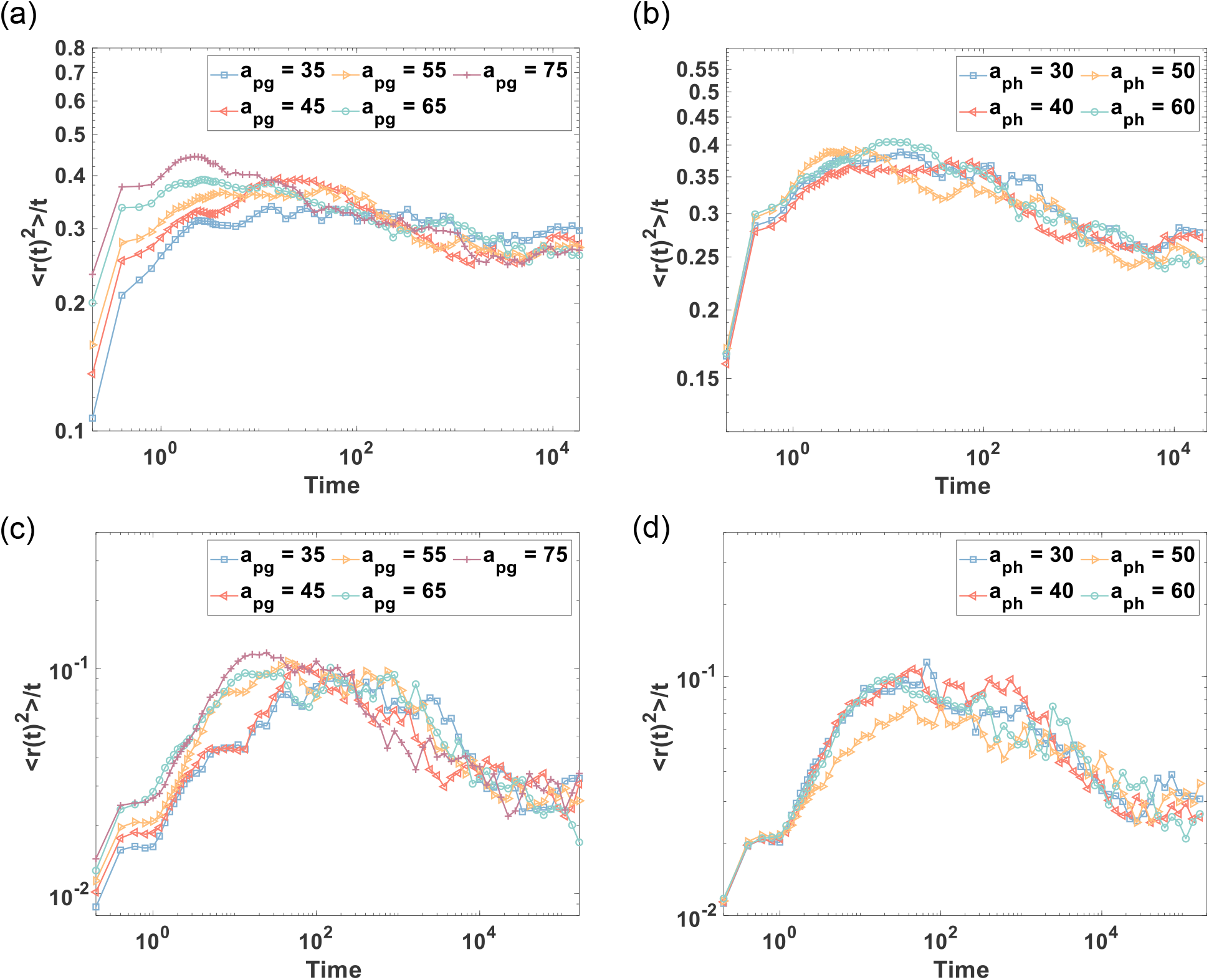
(a)-(b) The mean squared displacement of the small NPs (r = 0.5r_0_) with different PEG-Glycan repulsive parameters *a*_*pg*_ and PEG-Protein repulsive parameters *a*_*ph*_. (c)-(d) The mean squared displacement of the large NPs (r = 2r_0_) with different PEG-Glycan repulsive parameters *a*_*pg*_ and PEG-Protein repulsive parameters *a*_*ph*_.

**Figure 9:**
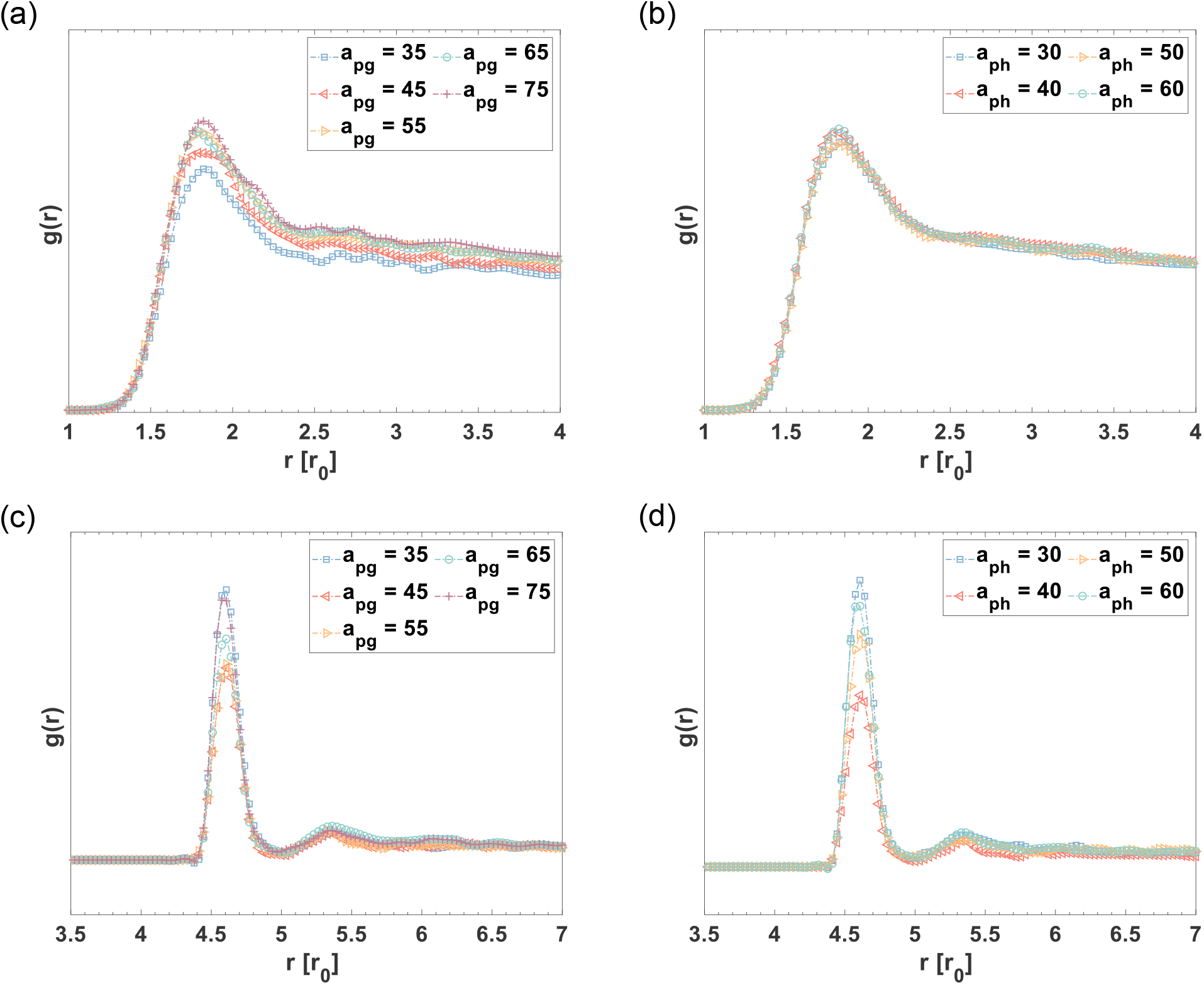
(a)-(b) The radial distribution function of the small NPs (r = 0.5r_0_) with different PEG-Glycan repulsive parameters *a*_*pg*_ and PEG-Protein repulsive parameters *a*_*ph*_. (c)-(d) The radial distribution function of the large NPs (r = 2r_0_) with different PEG-Glycan repulsive parameters *a*_*pg*_ and PEG-Protein repulsive parameters *a*_*ph*_.

Interestingly, we found that the degree of mucin cross-linking barely affected the diffusion rate of NPs, but significantly changed the mucin diffusion process (Figure 10 (a-b)). Especially for the mucin with 0% cross-linking, its MSD was very much higher than the cases of 50% and 100% cross-linking, indicating that the mucins were diffusing substantially, and its network could be altered to a large extent. To validate our observations, we calculated the pore size distributions of mucins after NPs diffusion (Figure 10 (c)). Compared with the pore size distribution of mucins without NPs (Figure 7 (a)) in all three cases, the addition of NPs severely disrupted the original normally-distributed pore sizes and led to the increase in pore sizes. However, the pore size distribution of the 0% cross-linked mucins was bimodal, having one peak centered around 2r_0_ and another peak around 7r_0_ to 8r_0_, showing that the mucins became denser at some regions but generated large gaps at other regions. On the other hand, we obtained the RDF of NPs to check how NPs aggregated in the three cases. The highest peak (blue line) indicates that NPs tended to aggregate more with themselves within the mucins with 0% cross-linking (Figure 10 (d)). All the data point to mucin-NPs phase separation occurring in mucin that lack cross-linking.

**Figure 10:**
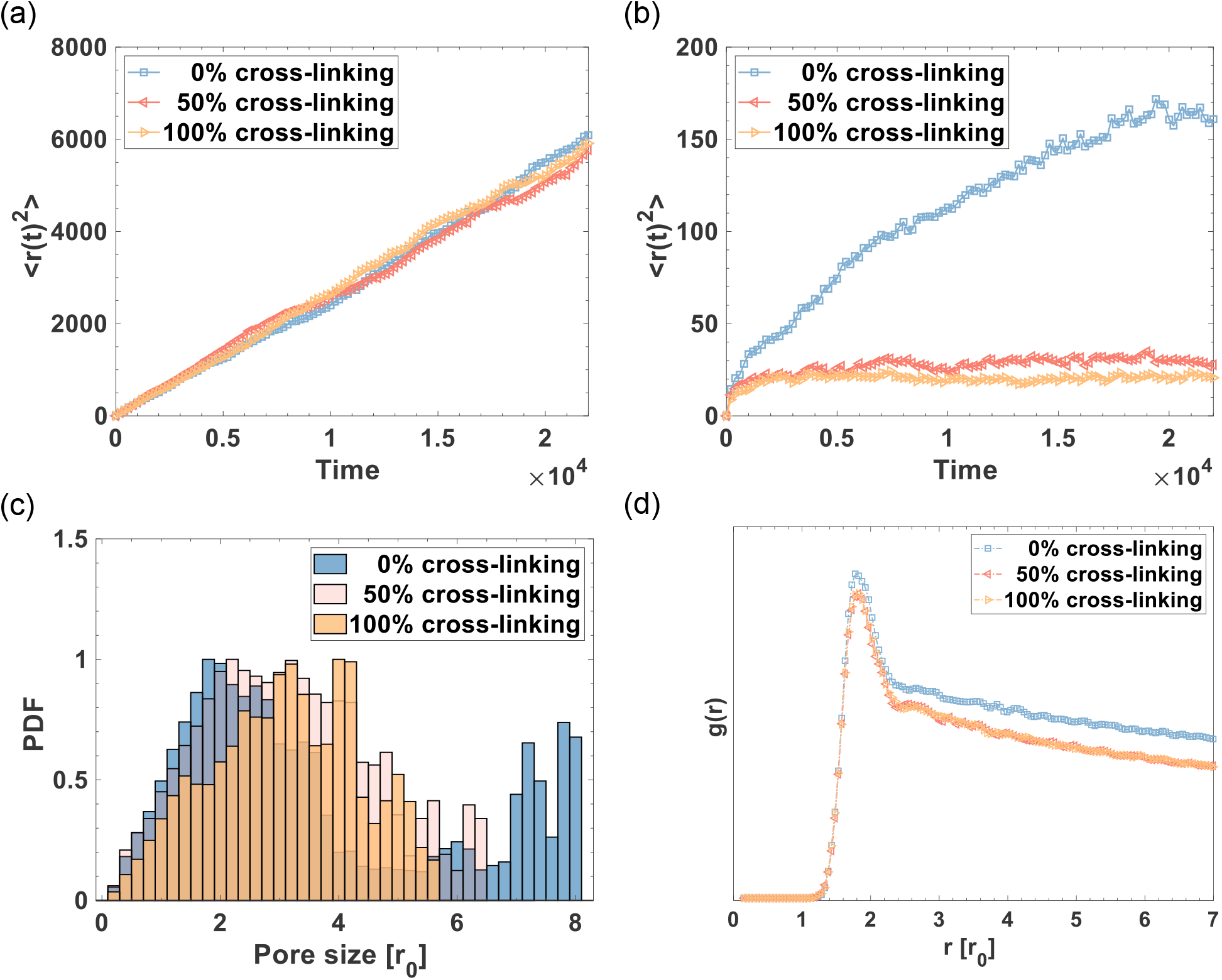
The MSDs of (a) NPs and (b) mucins under different degrees of cross-linking. (c) The pore size distributions of mucins with different degrees of cross-linking. (d) The radial distribution function of NPs.

## Conclusions

We developed a coarse-grained simulation model that specifically captures the molecular structures of mucus obtained directly from cultured human intestinal cells. To validate the model, we experimentally measured the rheological properties and the pore size of the mucus. Our model showed a good agreement with the experiment in terms of the steady shear viscosity. Furthermore, we studied the dynamics of nanoparticles diffusion inside the mucins based on the model. We first examined the influence of the NP size, finding that the diffusion coefficient of NPs was effectively reduced by the mucins, as the NPs size gradually increased to the pore size of mucins. Then, we found that the NPs-Mucins interactions mainly affected the ballistic regime rather than the Fickian diffusion regime, but it governed the NPs aggregations in the mucus network. Finally, we tested mucins with different degrees of cross-linking, and observed the phase separation phenomenon when NPs were in mucins with 0% cross-linking. While this model can be used to understand the diffusion of nanoparticle-based therapeutics, the model provides a promising platform for future work on the transport dynamics of microorganisms and drug particles in the human gastrointestinal mucus.

